# *Pseudomonas fluorescens 15* small RNA Pfs1 mediates transgenerational epigenetic inheritance of pathogen avoidance in *C. elegans* through the Ephrin receptor VAB-1

**DOI:** 10.1101/2024.05.23.595334

**Authors:** Renee Seto, Rachel Brown, Rachel Kaletsky, Lance R. Parsons, Rebecca S. Moore, Coleen T. Murphy

## Abstract

*C. elegans* are exposed to a variety of pathogenic and non-pathogenic bacteria species in their natural environment. Correspondingly, *C. elegans* has evolved an ability to discern between nutritive and infectious bacterial food sources. Here we show that *C. elegans* can learn to avoid the pathogenic bacteria *Pseudomonas fluorescens 15* (PF15), and that this learned avoidance behavior is passed on to progeny for four generations, as we previously demonstrated for *Pseudomonas aeruginosa* (PA14) and *Pseudomonas vranovensis*, using similar mechanisms, including the involvement of both the TGF-β ligand DAF-7 and *Cer1* retrotransposon-encoded virus-like particles. PF15 small RNAs are both necessary and sufficient to induce this transgenerational avoidance behavior. Unlike PA14 or *P. vranovensis*, PF15 does not use P11, Pv1, or a small RNA with *maco-1* homology for this avoidance; instead, an unrelated PF15 small RNA, Pfs1, that targets the *C. elegans vab-1* Ephrin receptor gene is necessary and sufficient for learned avoidance, suggesting the evolution of yet another bacterial sRNA/*C. elegans* gene target pair involved in transgenerational inheritance of pathogen avoidance. As VAB-2 Ephrin receptor ligand and MACO-1 knockdown also induce PF15 avoidance, we have begun to understand the genetic pathway involved in small RNA targeted pathogenic avoidance. Moreover, these data show that axon guidance pathway genes (VAB-1 and VAB-2) have previously unknown adult roles in regulating neuronal function. *C. elegans* may have evolved multiple bacterial specificity-encoded small RNA-dependent mechanisms to avoid different pathogenic bacteria species, thereby providing progeny with a survival advantage in a dynamic environment.

## Introduction

In their natural environment, *C. elegans* are exposed to a variety of bacterial food sources, some of which are pathogens. This microbe-rich environment has shaped *C. elegans* evolution, and ingestion of both nutritious and pathogenic bacteria can induce long-lasting molecular and behavioral changes in the worm. Our lab recently demonstrated that worms not only learn to avoid pathogenic PA14, but also pass on this learned avoidance behavior to their progeny for four generations^1^. Transgenerational epigenetic inheritance (TEI) is a mechanism by which parental experiences can be passed on to progeny, optimizing their responses for the environmental conditions experienced by their ancestors. Parental experiences and environmental factors lead to heritable changes in the gene expression and behavior of subsequent generations in a variety of organisms, including plants^2^, worms^3^, flies^4^, and mice^5,6^. In *C. elegans*, TEI has been demonstrated to play a role in a variety of processes, including antiviral^7^ and antibacterial immunity^8^, pathogen avoidance^1,3,9,10^, and other physiological responses to cellular stresses^11–14^.

Our lab previously found that *C. elegans* can pass on small RNA-mediated learned avoidance of *Pseudomonas aeruginosa* (PA14) or *Pseudomonas vranovensis* (GRb) to four generations of progeny through a molecular mechanism that requires an intact germline and neuronal signaling^1,3,9,10^. This process is initiated by the ingestion of bacterial small RNAs, and requires components of the RNA inheritance pathway, including SID-2, DCR-1, SID-1, and RNA-dependent RNA polymerases, as well as *Cer1* retrotransposon-encoded virus-like particles, and upregulation of the TGF-β ligand *daf-7* in the ASI sensory neurons^1,3,9,10^. Both *P. aeruginosa* P11 and *P. vranovensis* Pv1 sRNAs have homology to *C. elegans* gene *maco-1,* whose transcripts are downregulated in trained animals, resulting in upregulation of *daf-7* expression and subsequent avoidance behavior^3,10^. While P11 and Pv1 share no homology and target different exons of *maco-1*, their shared downregulation of *maco-1* explain why training on one bacteria causes avoidance of the other^10^.

Here, we show that transgenerational avoidance behavior can also be induced by another pathogenic bacteria strain, *Pseudomonas fluorescens 15* (PF15). PF15 was originally isolated from the rhizosphere of rice fields but belongs to a group of common soil bacterium that have been identified in the *C. elegans* microbiome^15,16^. Here, we show that transgenerational inheritance of PF15 avoidance lasts four generations after bacterial lawn training before preference returns to naïve levels. Furthermore, PF15 training induces increased *daf-7* expression in the ASI neuron pair, and *Cer1* virus-like particles are required for transgenerational learned avoidance, indicating that the previously-described molecular mechanisms of transgenerational epigenetic inheritance are used by PF15, as well. Uniquely, PF15 avoidance requires the small RNA *Pfs1* as a pathogen cue, which functions to downregulate the *C. elegans* gene *vab-1*, which encodes the Ephrin receptor. Knockdown of *vab-1* in adult *C. elegans*, like sRNA or Pfs1 treatment, induces avoidance of PF15; knockdown of *vab-2* (the Ephrin ligand) and *maco-1* also induce PF15 avoidance, pointing to the role of Ephrin receptor signaling and MACO-1 function in a shared pathway regulating pathogen avoidance. While the role of VAB-1 and VAB-2 have been shown in neuron development, these results suggest that the Ephrin signaling pathway has a novel adult function in regulation of pathogenic bacterial avoidance^17–19^. Thus, *C. elegans* appear to have evolved the ability to avoid several different pathogenic *Pseudomonas* species transgenerationally, but through independent examples of bacterial sRNA/*C. elegans* neuronal gene matches, eliciting species-specific avoidance responses.

## Results

### PF15 is pathogenic to *C. elegans* and can induce species-specific learned avoidance

*Pseudomonas fluorescens* (PF15) is pathogenic to *C. elegans*: after exposing worms to a lawn of PF15 for 24 hours, the worms become ill, their germlines shrink, and their lifespan is dramatically reduced (Figure 1A-C). Within 36 hours of PF15 exposure at 25 °C, over 80% of the worms have perished (median lifespan = 27hr; p<0.0001), while control (OP50-raised) worms will continue to live for an average of 2-3 weeks (Figure 1C). The innate immunity gene *irg-1* is induced upon exposure to PF15 (Figure 1D-E), as is also observed upon exposure to *P. aeruginosa* PA14 (but to a much lesser extent than in response to *P. vranovensis* GRb0427)^10^.

**Figure 1.**
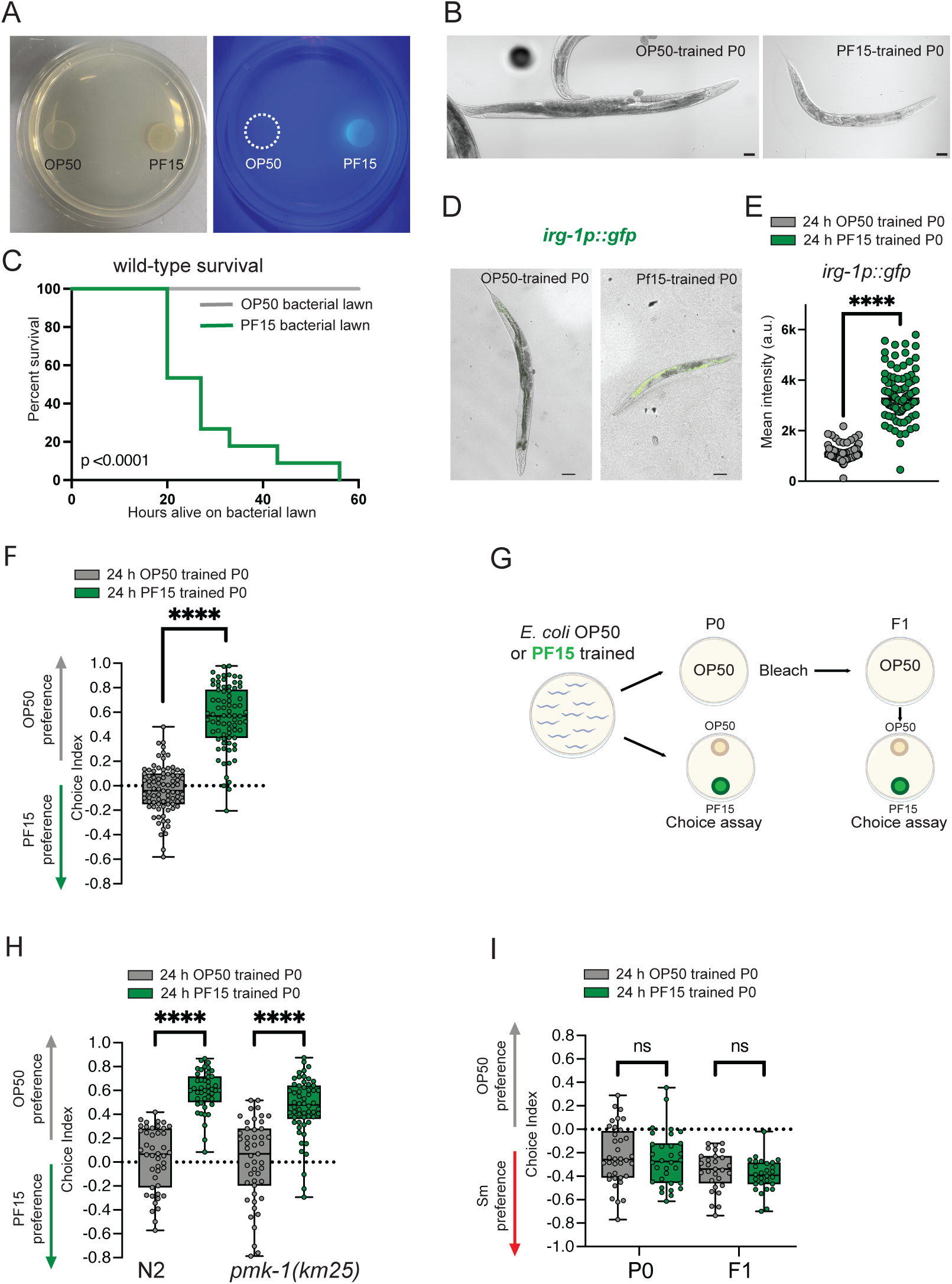
PF15 is pathogenic and induces species specific learned avoidance in *C. elegans*. (**A**) PF15 is fluorescent in UV light. Choice plates with spots of OP50 and PF15 bacteria. Left: image in white light. Right: image in UV light. (**B**) Representative images acquired after exposing Day 1 adults to OP50 (top) or PF15 (bottom) for 24 hours. PF15 is pathogenic and 24-hour exposure makes worms sick. Scale bar =100 μm. (**C**) *C. elegans* on PF15 lawn have reduced survival compared to survival on an OP50 lawn (25°C). (**D**) Representative images from one of three biological replicates of *irg-1p::gfp* expression in Day 1 adults exposed to OP50 (left) or PF15 (right) for 24h starting at L4. Images are merged brightfield and GFP channels. Scale bar = 100 μm. (**E**) Quantification of *irg-1p::gfp* in P0 PF15-trained worms. Expression of *irg-1p::gfp* is induced by PF15 bacterial lawn exposure. Representative of three biological replicates. (**F**) Adult *C. elegans* trained for 24h on PF15 induces PF15 avoidance compared to OP50-trained control. (**G**) Adult *C. elegans* pathogen training protocol and choice assay. After training, worms are used to either test preference in a choice assay or bleached to obtain F1 progeny. Choice Index = (# of worms on OP50-# of worms on PF15)/(total # worms). (**H**) *pmk-1(km25)* mutants can learn PF15 avoidance from PF15 lawn training. (**I**) *C. elegans* trained on PF15 for 24 h do not learn to avoid pathogen *Serratia marcescens* (*S. marcescens*). F1 progeny of PF15-trained worms also do not avoid *S. marcescens.* Each dot represents an individual worm (E) or an individual choice assay plate (F, H-I). Boxplots: center line, median; box range, 25th–75th percentiles; whiskers denote minimum-maximum values. Unpaired, two-tailed Student’s t test, ****p < 0.0001 (E-F); one-way ANOVA with Tukey’s multiple comparison’s test, ns, not significant (I); Two-way ANOVA with Tukey’s multiple comparison’s test, ****p < 0.0001 (H). For the survival assay in (C), ****p<0.0001 (by Log-rank (Mantel-Cox) test for survival).

Despite its pathogenicity, naive *C. elegans* show a slight naïve preference to PF15 over the lab strain of nonpathogenic *E. coli* (OP50) (Figure 1F). However, after 24 hours of exposure to a PF15 lawn, *C. elegans* robustly avoid PF15 (Figure 1F, G). The innate immunity regulator *pmk-1* is not required for avoidance after training on PF15 bacterial plates (Figure 1H), suggesting that other pathways may be functional in P0 avoidance besides innate immunity-induced avoidance, as has previously been shown^20–23^. Additionally, PF15 training does not induce avoidance of another bacterial pathogen, *Serratia marcescens (S. marcescens)*, in P0 or F1, indicating that this is not a universal pathogen response (Figure 1I).

### PF15 induces transgenerational epigenetic inheritance of avoidance

Like *P. aeruginosa* (PA14)- and *P. vranovensis* (GRb0427)-induced transgenerational epigenetic inheritance (TEI) of avoidance behavior, 24hr of exposure to PF15 also induces *C. elegans’* TEI of pathogen avoidance behavior (Figure 2A): this learned avoidance behavior in P0 mothers was then passed on to progeny for four generations without any additional training, while fifth generation descendants resume naïve behavior, no longer avoiding PF15 (Figure 2B). As we previously observed with PA14 training, the magnitude of P0 avoidance is greater than subsequent generations, likely due to the combined effect of innate and transgenerational immune responses.

**Figure 2.**
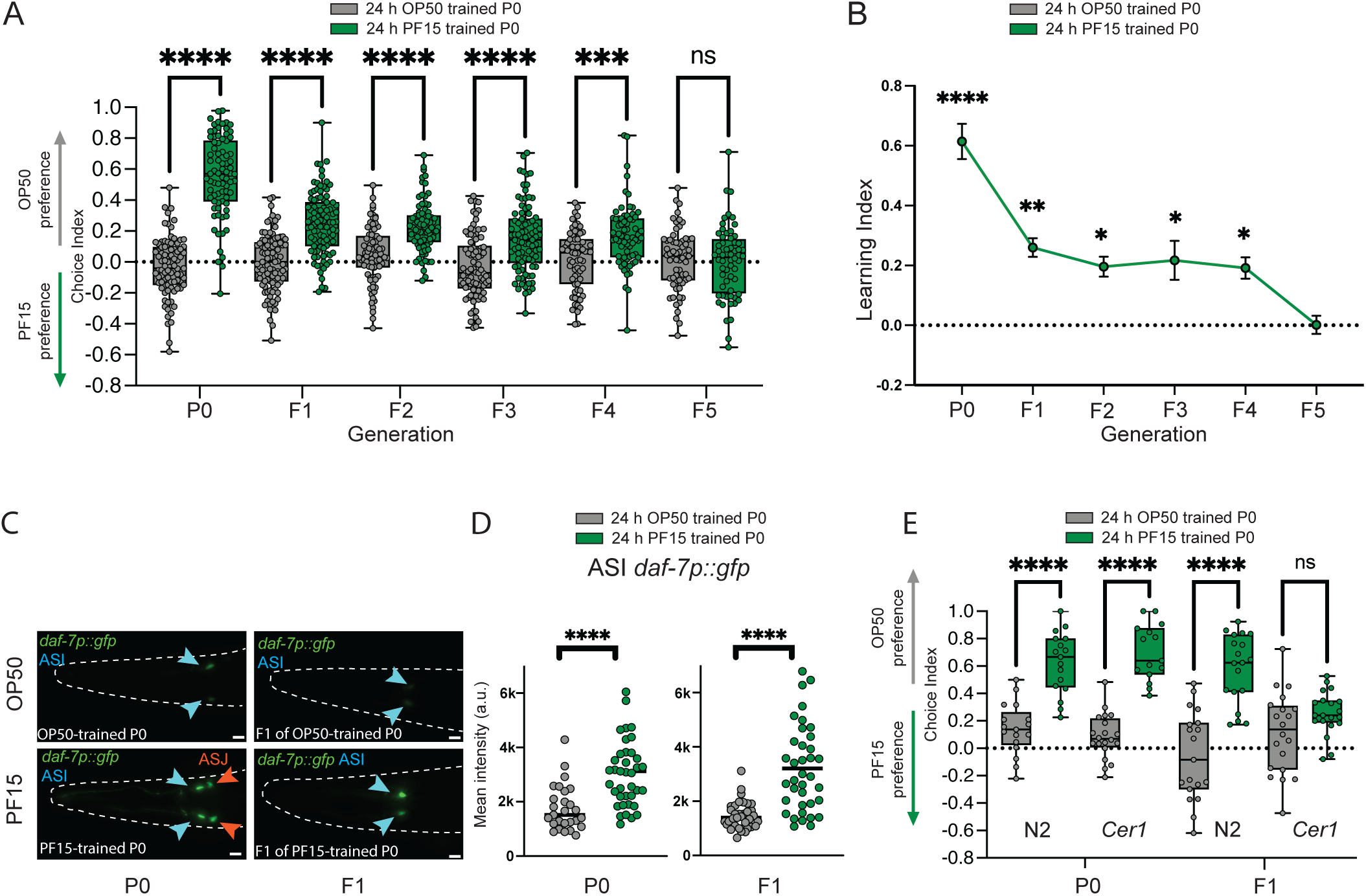
PF15 exposure induces transgenerational epigenetic inheritance of PF15 avoidance and requires conserved components of sRNA-mediated transgenerational avoidance pathway. (**A**) Transgenerational inheritance of pathogen avoidance: P0 worms trained on a PF15 bacterial lawn for 24h learn to avoid PF15 compared to OP50 control, and their F1-F4 progeny retain PF15 avoidance behavior until it returns to baseline in F5. (**B**) Learning index (naive choice index – trained choice index) of generation P0–F5. Error bars represent mean ± SEM. (**C**) Representative images of 24hr OP50-trained or PF15-trained worm expressing *daf-7p::gfp;* fluorescence is increased in the ASI (blue arrowheads) and ASJ sensory neurons (orange arrowhead) in P0. P0 worms (left) trained on PF15 (bottom) show increased *daf-7p::gfp* in the ASI and ASJ compared to OP50-trained controls (top). F1 progeny (right) of PF15-trained P0 (bottom) also show increased *daf-7p::gfp* in the ASI only compared to F1 progeny of OP50-trained P0 (top). Scale bar = 10 μm. (**D**) Quantification of *daf-7p::gfp* in P0 24h OP50- and PF15-trained worms and their F1 progeny. PF15 training increases *daf-7p::gfp* expression in the ASI neuron pair in P0 and F1. P0 representative of one of three biological replicates. (**E**) *Cer1(gk870313)* mutants learn PF15 avoidance in P0 but are unable to avoid PF15 in F1. Each dot represents an individual neuron (D) or an individual choice assay plate (A, E). Boxplots: center line, median; box range, 25th–75th percentiles; whiskers denote minimum-maximum values. Unpaired, two-tailed Student’s t test, ****p < 0.0001 (D); one-way ANOVA with Tukey’s multiple comparison’s test, ****p<0.0001, ***p<0.001, **p<0.01, *p>0.05, ns, not significant (A, B); Two-way ANOVA with Tukey’s multiple comparison’s test, ****p < 0.0001, ns, not significant (E).

### DAF-7/TGF-β expression is required for transgenerational inheritance of PF15 avoidance

Neuronal function is required for the sensing and subsequent avoidance of pathogenic bacteria strains. With PA14 or *P. vranovensis* exposure, neuronal expression of the TGF-β ligand *daf-7* is dramatically altered: in mothers exposed to PA14, *daf-7* expression is induced in the ASI and ASJ neurons, and their progeny exhibit increased *daf-7* expression specifically in the ASI neurons. PA14 sRNA, P11, GRb sRNA, or Pv1 sRNA exposure also induce *daf-7p::GFP* fluorescence in the ASI in P0 through F4 generations^1,3,10,20^, indicating that the ASI expression of *daf-7* is indicative of sRNA-mediated avoidance. We were curious whether *daf-7* signaling also correlates with inheritance of PF15 avoidance. Worms exposed to a lawn of PF15, like PA14, show increased *daf-7p::GFP* expression in the both the ASI and the ASJ neurons in the P0 generations (Figure 2C), and remains elevated in the F1 generation, as we previously found for PA14 (Figure 2C, D). Although PF15 does not exhibit the blue color that is the hallmark of PA14 phenazine production, PF15 does produce pyoverdine, a fluorescent siderophore (Figure 1A)^24,25^. F1 progeny who inherit PF15 avoidance behavior maintain high ASI *daf-7p::GFP* expression levels, which correlates with avoidance (Figure 2A)^1,3,10^.

### *Cer1* is required for transgenerational inheritance of PF15 avoidance

The TEI of PA14 avoidance requires *Cer1* retrotransposon-encoded virus-like particles, which act at the step of transfer information from the germline to neurons, upstream of *daf-7* regulation^9^. We were curious whether the TEI of PF15 avoidance also requires *Cer1*, or if PF15 avoidance utilizes another mechanism for germline-to-neuron signaling. We found that learned avoidance of PF15 is not inherited by the F1 generation in *Cer1(gk870313)* mutants (Figure 2E). Thus, these results suggest that *Cer1* is required for the TEI of PF15 avoidance, as it is for PA14- and *P. vranovensis*-induced avoidance^9,10^.

### PF15-derived sRNAs induce learned pathogen avoidance and TEI behavior without illness

Since purified sRNA isolated from PA14 is both sufficient and necessary to induce transgenerational avoidance of PA14 in worms^3^, and we observe similar mechanisms and F1-F4 TEI of learned PF15 avoidance, we were curious whether PF15 sRNAs also regulate the transgenerational avoidance of PF15. To test this hypothesis, we first isolated sRNAs (<200 nt) from PF15 or OP50 (control) lawns (see Methods). We then trained worms by adding these purified small RNAs to an OP50 *E. coli* lawn for 24hrs. Although worms trained on sRNA-treated OP50 for 24hr are healthy (Figure 3A), PF15-derived sRNA training was sufficient to induce learned avoidance behavior in the trained worms (Figure 3B), and this avoidance lasts for four additional generations (F1-F4; Figure 3C). The magnitude of the small RNA-dependent response in P0 animals is smaller than the lawn training P0 response, consistent with an additive effect of innate immune response and small RNA response in the P0 generation (Figure 3D) that we have similarly observed with PA14^3^. Furthermore, *Cer1(gk870313)* worms trained with PF15 sRNA are unable to learn to avoid PF15 (Figure 3E), consistent with the previously-determined role for *Cer1* in transmitting sRNA-based information from the germline to neurons, even in the P0 generation. Together, these results suggest that the mechanism for PF15 sRNA-mediated avoidance in mothers and their progeny is similar that observed for PA14-induced avoidance^1,3,9^. The fact that purified sRNAs are sufficient to induce avoidance of PF15 in the absence of intact pathogen suggests that PF15 sRNA-induced avoidance does not require virulence. Thus, sRNA-mediated learned avoidance functions independently of the innate immune response.

**Figure 3.**
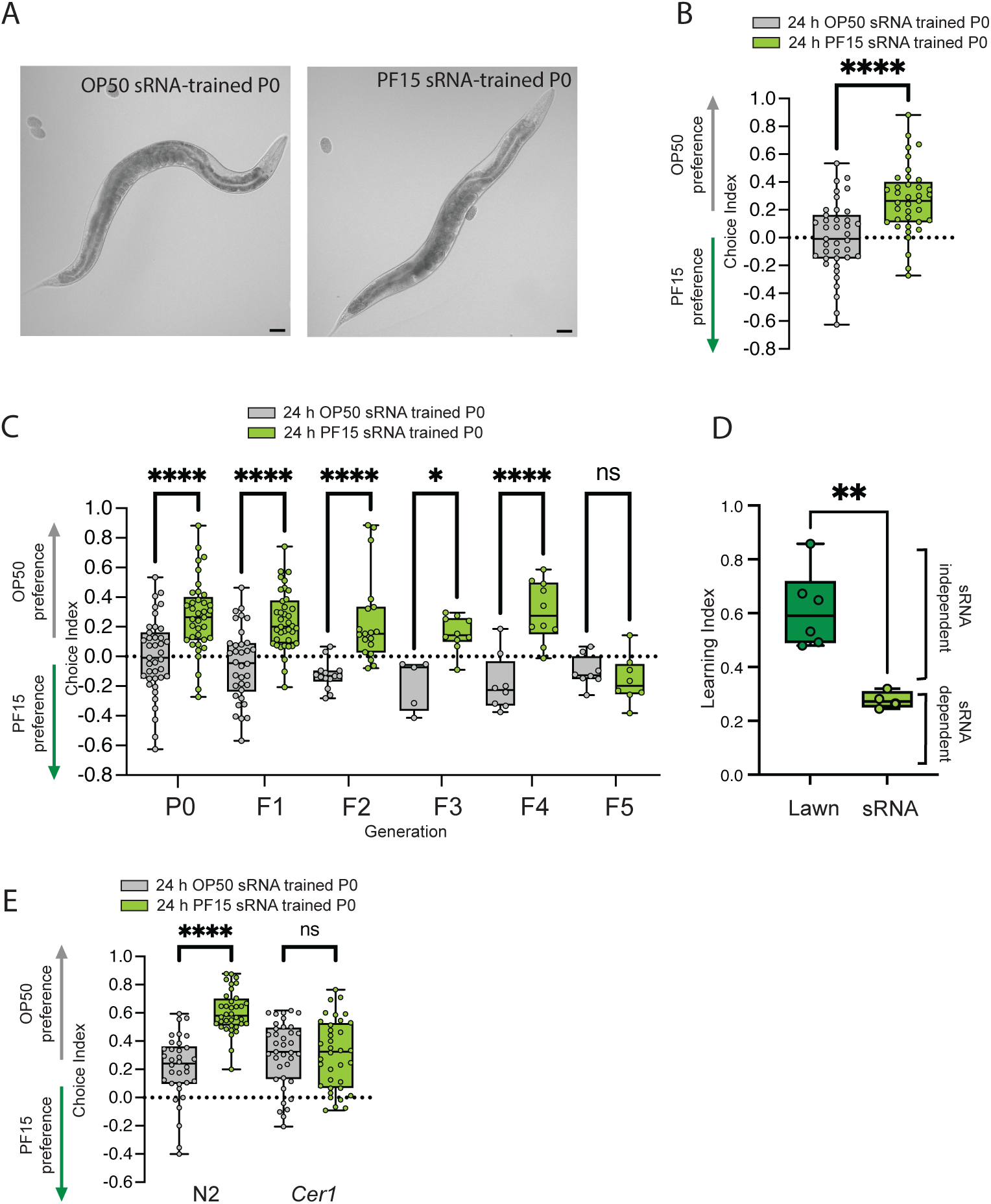
PF15 sRNAs are sufficient to cause transgenerational PF15 avoidance and do not activate the innate immune system. (**A**) Representative images of Day 1 worms on OP50 sRNAs (left) or PF15 sRNAs (right) added to OP50 lawns for 24h. Worms treated with PF15 sRNAs do not appear sick, unlike lawn trained animals (see Fig. 1B). Scale bar = 50 μm. (**B**) Training with PF15 sRNAs for 24h causes PF15 avoidance compared to OP50 sRNA-trained control. (**C**) Transgenerational inheritance of pathogen avoidance: P0 worms trained on PF15 sRNAs for 24h learn to avoid PF15 compared to OP50 control, and their F1-F4 progeny retain PF15 avoidance behavior until it returns to baseline in F5. (**D**) Comparative magnitude of PF15 lawn vs PF15 sRNA-only P0 avoidance suggests that P0 avoidance is the cumulative effect of sRNA-dependent and sRNA-independent avoidance pathways. (**E**) *Cer1* mutants are unable to learn PF15 avoidance from PF15 sRNA training in P0 or F1. Each dot represents an individual choice assay plate (B-C, E). Boxplots: center line, median; box range, 25th–75th percentiles; whiskers denote minimum-maximum values. Unpaired, two-tailed Student’s t test, ****p < 0.0001, **p>0.01 (B, D); one-way ANOVA with Tukey’s multiple comparison’s test, ****p<0.0001, *p>0.05, ns, not significant (C); Two-way ANOVA with Tukey’s multiple comparison’s test, ****p < 0.0001, ns, not significant (E).

### Identification of PF15 sRNAs required for PF15 avoidance

Our results suggested that PF15-induced TEI of avoidance is mediated by small RNAs, but the specific sRNA was unknown. Because there was no publicly available PF15 genome sequence that we could mine for information, we first purified, sequenced, and annotated the PF15 genome (Figure 4A; BioProject PRJNA1114673; see Methods). Because the simplest model would be that PF15 would contain either the same small RNA as we previously identified in other *Pseudomonas* strains, we BLASTed the P11 sequences against the PF15 genome, but we found no significant sequence similarity to either. Next, reasoning that P11 targets the *C. elegans maco-1* gene with a 17nt stretch of perfect match to exon 8 of *maco-1*^3^, we also BLASTed the sequence of *maco-1* against the PF15 genome, but found no hits within intergenic regions, where small RNAs are usually found^26,27^. This was in contrast to *P. vranovensis*, which encodes a small RNA, Pv1, in an intergenic region; Pv1 also has no similarity to P11 but does have a 16nt match to exon 1 of *maco-1*^10^. Together, these results suggest that P11 is not the small RNA responsible for inducing TEI of PF15 avoidance, and that the relevant PF15 small RNA targets a *C. elegans* gene other than *maco-1*.

**Figure 4.**
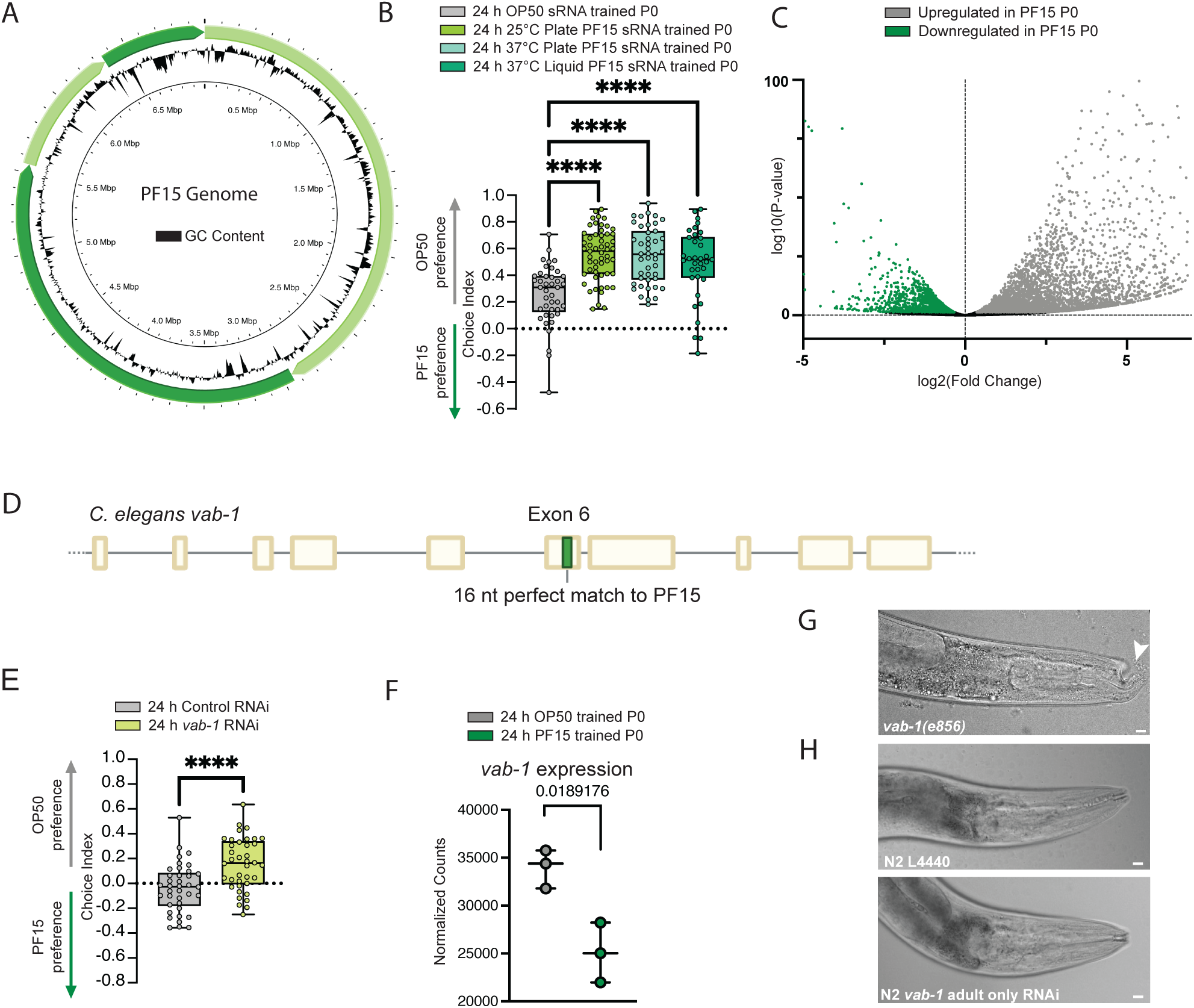
Sequencing the PF15 genome and PF15 sRNAs identifies the *C. elegans* gene *vab-1* as a regulator of PF15 avoidance. (**A**) Circular representation of the PF15 genome. Outer circle: green segments represent individual contigs. Middle circle: GC content is indicated. Inner circle: Genome length is indicated. Generated by Proksee. (**B**) sRNAs were collected from PF15 in three growth conditions: 25°C plate, 37°C plate, and 37°C liquid, and used for training alongside OP50 sRNA control. PF15 sRNAs from all three conditions cause PF15 avoidance. (**C**) Differential mRNA expression of P0 *C. elegans* trained on OP50 vs. PF15 lawn for 24 h. (**D**) The PF15 genome contains a 16 nt perfect match to the *vab-1* gene in exon 6. (**E**) Worms exposed to adult only *vab-1* RNAi naively avoid PF15 compared to control. (**F**) *vab-1* expression is decreased in PF15-trained P0 worms, as shown by differential mRNA expression. Adjusted p-value= 0.0189176. (**G**) The *vab-1(g995)* mutant exhibits gross morphological defects in the head. Scale bar = 100 μm. (**H**) Image of N2 worm treated with adult-only RNAi (24h, L4 to Day 1) of *vab-1* shows no morphological defects. Scale bar = 100 μm. Each dot represents an individual choice assay plate (B, E) or a sequencing replicate (F). Boxplots: center line, median; box range, 25th–75th percentiles; whiskers denote minimum-maximum values. Unpaired, two-tailed Student’s t test, ****p < 0.0001, (D); one-way ANOVA with Tukey’s multiple comparison’s test, ****p<0.0001 (B).

To narrow our search for small RNAs that might target *C. elegans* genes, we then identified the intergenic regions in PF15 through gene prediction, indicating possible locations of sRNAs, as we had found for Pv1 in the *Pseudomonas vranovensis* genome^26,27^. Of these intergenic regions, we then determined which had ≥16 nucleotides of perfect sequence match to *C. elegans* genes, resulting in a list of 11,423 intergenic regions and their corresponding *C. elegans* genes.

To further narrow down which small RNA might be responsible for PF15’s induction of transgenerational avoidance, we tested PF15 grown under three different conditions. We found that small RNAs isolated from 25°C plate (our usual growth condition), 37°C plate, and 37°C liquid growth conditions all induced avoidance (Figure 4B). These results contrast with PA14, which only induces avoidance when bacteria are grown on plates at 25°C, allowing differential expression analysis to narrow down the set of possible candidates to only six sRNAs^3^. Because all three conditions induced avoidance, we then examined the sRNAs in the intersection between the three conditions, narrowing the list to 646 candidate sRNAs.

To identify candidate target *C. elegans* genes, we isolated and compared mRNA from worms that had been trained on PF15 or OP50 control bacteria (Figure 4C). Because previous target *C. elegans* gene *maco-1* transcripts are downregulated in worms treated with PA14 or *P. vranovensis*^10^, we examined the list of genes that were downregulated upon PF15 treatment, resulting in a list of 174 candidates. Then we noted which of these candidates into those that were also downregulated in PA14 treated worms, and those that overlap. These filtering steps resulted in a candidate list of 26 candidate intergenic regions that correspond to *C. elegans* genes.

Previously, we found that mutants or RNAi knockdown of *maco-1* increased naïve avoidance of PA14^3^. Therefore, to test whether any of these genes might be targets of PF15 small RNAs, we carried out a rapid candidate screen by measuring the naive avoidance of worms with a candidate gene knocked down in adulthood; while 18 of the 19 available RNAi candidate gene knockdowns had no effect, reduction of *vab-1,* which has a 16nt match with an intergenic region of the PF15 genome (Figure 4D), significantly affected naïve attraction to PF15. We repeated this adult-only RNAi knockdown of *vab-1* and found that reduction of *vab-1* does indeed increase naïve avoidance of PF15 (Figure 4E), similar to the high naïve avoidance of PA14 by *maco-1* RNAi and *maco-1* mutants^3^. We also confirmed that *vab-1* gene expression is significantly decreased in worms treated with PF15 (Figure 4F).

To further test the possible role of *vab-1* in PF15 avoidance, we obtained genetic loss-of-function mutants of *vab-1* (*vab-1(e856)*); however, these animals exhibit significant developmental defects and are too defective to test (Figure 4G). By contrast, adult-only RNAi of *vab-1* allows normal development (Figure 4H) and chemotaxis (Figure 4E), underscoring the advantage of using RNAi in adulthood to unmask phenotypes that might never be found using mutants that have developmental phenotypes.

### Pfs1 sRNA causes transgenerational PF15 avoidance and contains a match to *C. elegans* gene *vab-1*

To test if the PF15 intergenic region we identified caused avoidance, we cloned the 600 bp intergenic fragment containing the *vab-1* homology region into *E. coli* and used this bacteria to train worms, as previously described^3,10^. We found that worms trained on the intergenic region *E. coli* learned to avoid PF15 from P0 through F4 (Figure 5A).

**Figure 5.**
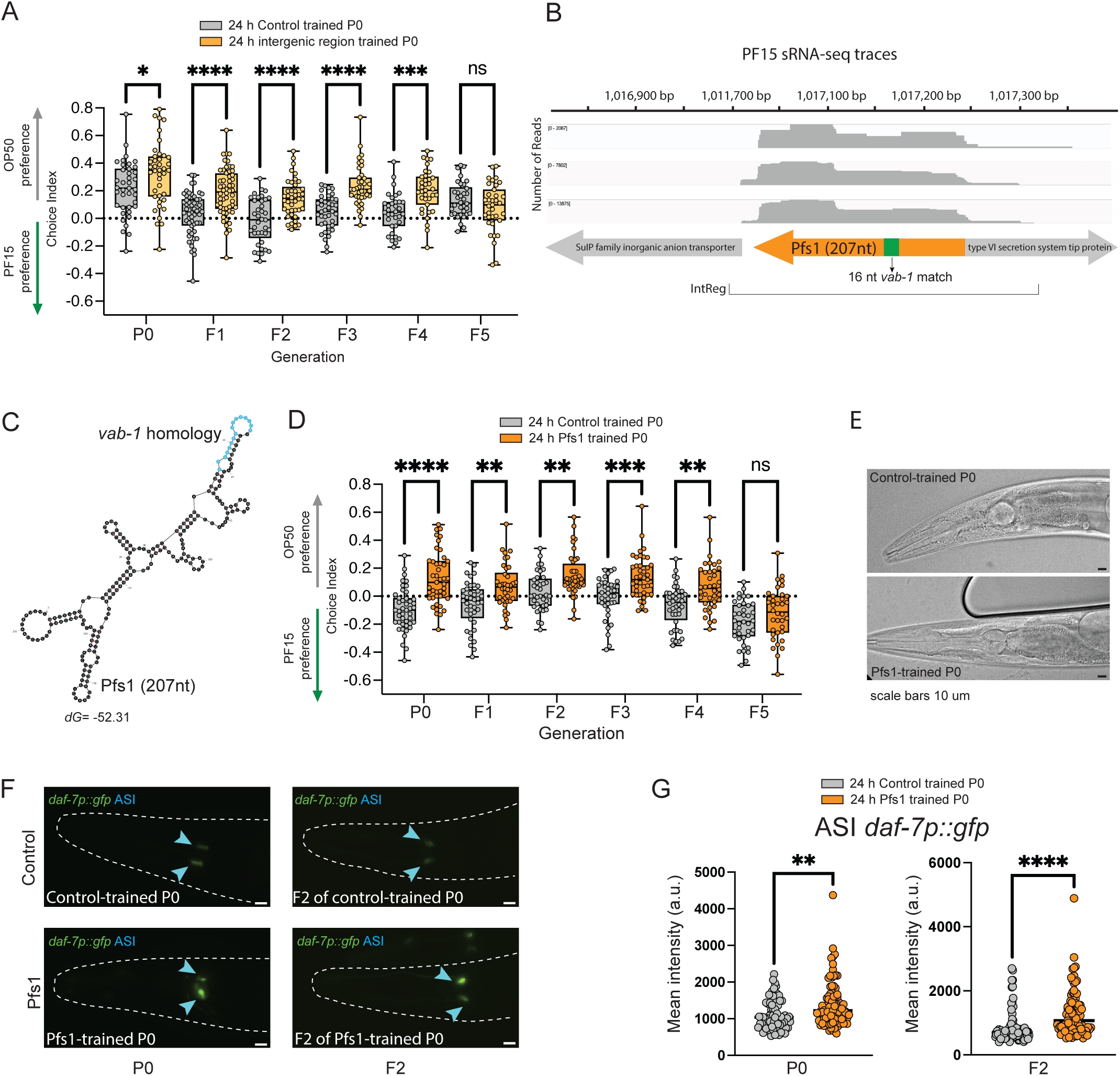
Training with Pfs1 sRNA causes transgenerational avoidance of PF15. (**A**) Training with the PF15 intergenic region containing the *vab-1* homology causes PF15 transgenerational avoidance from P0 through F4, and resumes normal naïve behavior in F5. (**B**) Reads from sRNA sequencing reveal a potential sRNA in the intergenic region we had identified. This putative sRNA, Pfs1, contains a 16 nt perfect match to the *C. elegans* gene *vab-1*. The intergenic region containing the Pfs1 small RNA is between two genes, a SuIP family inorganic anion transporter and a type VI secretion system tip protein. The region cloned and used for (A) is indicated. (**C**) Rapid amplification of cDNA ends identified an sRNA of 207 nt in length. Pfs1 sRNA secondary structure predicts that the region containing the 16nt match to the *C. elegans* gene *vab-1* is in a hairpin structure. (**D**) Training with the Pfs1 sRNA results in transgenerational PF15 avoidance from P0-F4. (**E**) Pfs1 adult-only trained animals (P0) show no morphological defects. Scale bar = 100 μm. (**F**) Representative images of 24hr control-trained or Pfs1-trained *daf-7p::gfp* worms; fluorescence is increased in the ASI sensory neurons (blue arrowheads) of Pfs1-trained relative to control 24h trained worms (top), and is maintained in F2 grandprogeny Scale bar = 100 μm. Representative of three biological replicates. (**G**) Quantification of *daf-7p::gfp* in P0 24h control and Pfs1 sRNA-trained worms and their F2 grandprogeny. PF15 training increased *daf-7p::gfp* expression in the ASI neuron pair in P0 and F2. Representative of three biological replicates. Each dot represents an individual choice assay plate (A, D) or an individual neuron (G). Boxplots: center line, median; box range, 25th–75th percentiles; whiskers denote minimum-maximum values. Unpaired, two-tailed Student’s t test, ****p < 0.0001, **p<0.01 (G); one-way ANOVA with Tukey’s multiple comparison’s test, ****p<0.0001, ***p<0.001, **p<0.01, *p<0.05 (A, D).

To identify the small RNA with *vab-1* homology expressed within the intergenic region, we examined our sRNA sequencing data (Figure 4) and observed that there are reads covering the region between a SuIP family inorganic anion transporter and a type VI secretion system tip protein, with one sRNA common to all conditions of ∼200 nt that includes the 16 nt of sense *vab-1* homology, which we named Pfs1 (Figure 5B). To precisely define the ends of the Pfs1, we used rapid amplification of cDNA ends (RACE) to sequence the 5’ and 3’ ends of this small RNA; RACE results suggest that Pfs1 is 207 nucleotides long (Figure 5B). Once the precise ends of the sRNA had been defined, we generated predicted secondary structures for the Pfs1 sRNA. Although there are several possibilities for these predicted structures, several of the models predicted that the 16nt match lies in a stem-loop, as was previously predicted for the P11 and Pv1 structures (Figure 5C).

We then tested the Pfs1 small RNA by cloning it into *E. coli*, then training *C. elegans* on this *E. coli* expressing Pfs1, which induces avoidance of PF15 from P0-F4 before returning to naïve preference in F5, as we had seen with our lawn and total small RNA training (Figure 3). Once again, we note that training on the sRNA alone (Pfs1) induces the same level of avoidance in P0 as in F1-F4 (Figure 5D), suggesting that no additional innate immunity effect is functioning in the P0. Despite reducing *vab-1* levels (Figure 5E), Pfs1-trained animals (Figure 5F) do not exhibit the morphological defects of the *vab-1* mutants consistent with adult-only *vab-1* RNAi training.

We next wondered if Pfs1 training was responsible for the increase in ASI *daf-7p::gfp* levels we had observed in PF15 lawn and total sRNA training. Indeed, we found that training worms on Pfs1-expressing bacteria for 24h significantly increased ASI *daf-7p::gfp* levels compared to control-trained worms, and the F2 progeny of trained P0 mothers retained this pattern (Figure 5G, H). Therefore, exposure to the small RNA Pfs1 is sufficient to induce transgenerational avoidance of PF15 and *daf-7p::gfp* expression.

### vab-1 acts upstream of maco-1

We wondered whether *vab-1* acts in the same or different pathway as *maco-1*, the gene targeted by both PA14’s P11 sRNA and *P. vranovensis’* Pv1 sRNA. First, we tested whether *maco-1* mutants display a high naïve avoidance of PF15, as they do to PA14; in fact, they do (Figure 6A), despite the fact that the target of PF15 small RNA is *vab-1*, not *maco-1*. Similarly, *maco-1* RNAi also induces naïve avoidance of PF15 (Figure 6B). We then asked whether maco-1’s high naïve avoidance of PF15 is increased by Pfs-1 treatment; however, *maco-1* mutants’ high naïve avoidance is not affected by Pfs1 treatment (Figure 6C). Together, these data suggest that *vab-1* and *maco-1* act in the same pathway to control pathogen avoidance.

**Figure 6.**
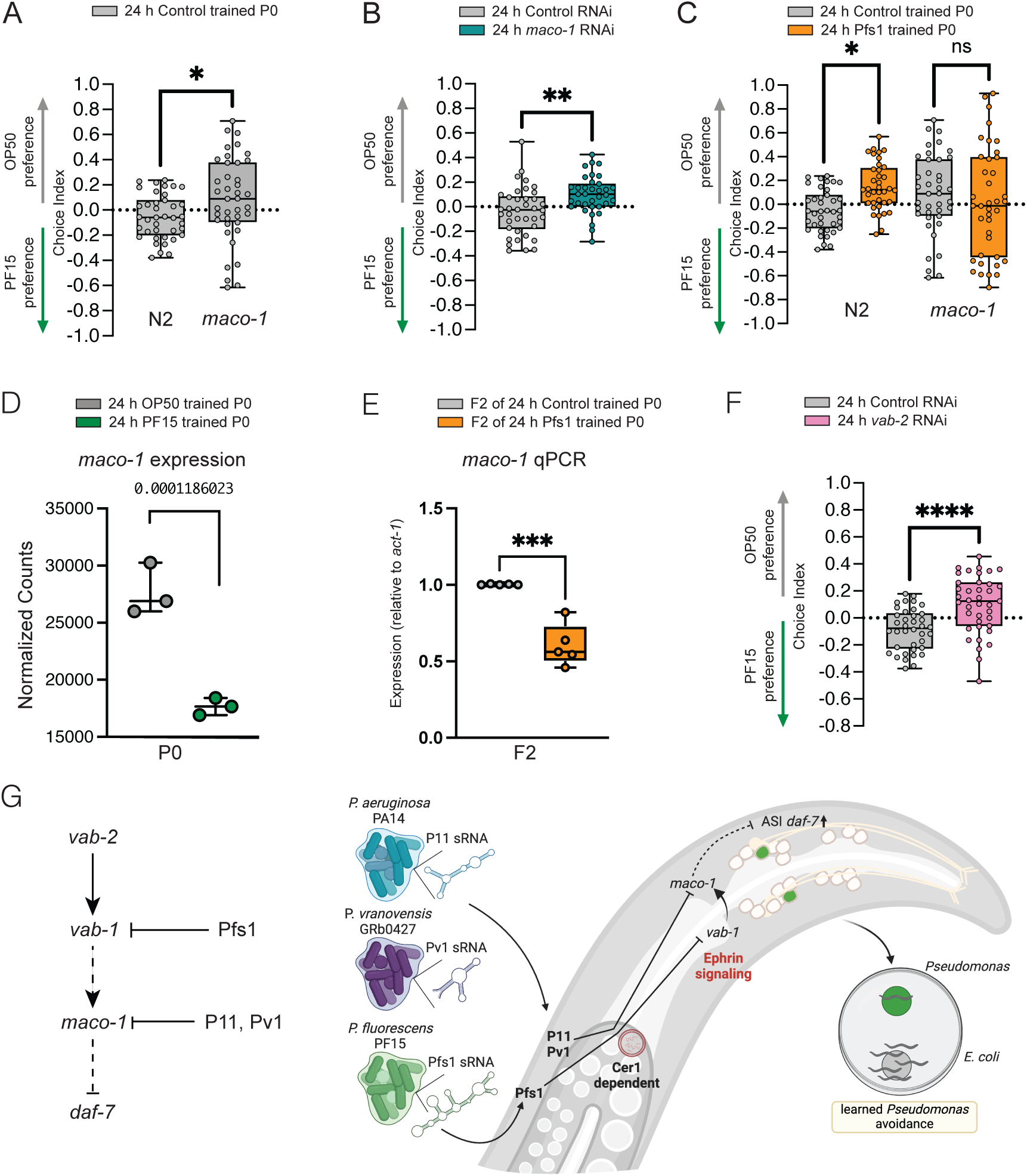
*vab-1* acts upstream of *maco-1* to induce PF15 avoidance. (**A**) *maco-1(ok3165)* loss-of-function mutant worms naively avoid PF15 relative to wild type worms. (**B**) Worms exposed to adult-only *maco-1* RNAi naively avoid PF15 compared to control. (**C**) *maco-1(ok3165)* loss-of-function mutant worms do not show increased PF15 avoidance learning with Pfs1 training. (**D**) *maco-1* expression is decreased in PF15-trained P0 worms, as shown by differential mRNA expression. Adjusted p value= 0.0001186023. (**E**) qPCR of *maco-1* levels in F2 grandprogeny of Pfs1-trained animals. Fold change (2^(-ΔΔCt) of *maco-1* transcript levels in Pfs1-trained F2 animals relative to the respective OP50-trained control (*act-1* was used as the housekeeping gene for reference). Each data point represents an independent biological replicate, and 3 technical replicates were performed for each biological replicate. (**F**) Worms exposed to adult-only *vab-2* RNAi naively avoid PF15 compared to control. (**G**) Model of *vab-1* epistasis and mechanism of Pfs1-mediated PF15 avoidance in comparison to previous findings of P11 and Pv1 mediated avoidance^3,10^. Each dot represents an individual choice assay plate (A-C, F) or a biological replicate (D, E). Boxplots: center line, median; box range, 25th–75th percentiles; whiskers denote minimum-maximum values. Unpaired, two-tailed Student’s t test, ****p < 0.0001, **p<0.01, *p>0.05, ns, not significant (A-B, E-F); Two-way ANOVA with Tukey’s multiple comparison’s test, *p < 0.05, ns, not significant (E).

While *vab-1* and *maco-1* acting in the same pathway might be somewhat expected, our original model did not suggest that they might have an effect on one another. Surprisingly, however, our transcriptional analyses show that *maco-1* is downregulated in PF15-treated worms (Figure 6D), and qPCR of F2 (grandprogeny) of Pfs1-treated animals show that *maco-1* transcripts are downregulated transgenerationally (Figure 6E). That is, despite neither PF15 nor Pfs1 having any match to *maco-1* that might suggest an RNAi-like mechanism at work, *maco-1* transcripts are decreased, perhaps suggesting that decreased VAB-1 signaling might affect the transcription of the *maco-1* gene.

### The VAB-1 ephrin receptor and its ligand, VAB-2 are required for PF15 avoidance

*vab-1* encodes the only known ephrin receptor in *C. elegans*. *vab-1* is expressed in many tissues of the developing larva (hypodermal cells, neuroblasts, neurons, and oocytes), is required in development for coordinated cell movements, and is required for normal epidermal morphogenesis; *vab-1* mutants have severe morphological defects (Figure 4F)^17–19,28^. In adulthood, *vab-1* is expressed in adult neurons^29^ as well as other tissues, but has no previously reported functions in adults. We wanted to know if any characterized upstream components of the *vab-1* pathway were required for PF15 avoidance. To test this, we treated adult N2 worms with RNAi against the worm ephrin *vab-2* and found that these worms avoided PF15 (Figure 6F), suggesting that the VAB-2 Ephrin ligand and its receptor play a role in the avoidance choice. Together, these data suggest that *vab-2, vab-1*, and *maco-1* all function in the same pathway to regulate avoidance of pathogenic *Pseudomonas* species through bacterial sRNA targeting.

## Discussion

Previously, we found that both *Pseudomonas aeruginosa* (PA14) and *Pseudomonas vranovensis* (GRb0427) encode small RNAs with short perfect matches to the *C. elegans* gene *maco-1* that induce avoidance of those pathogens in the mother and four additional generations^1,3,9,10^. Here we show that yet a third *Pseudomonas* species, *Pseudomonas fluorescens* PF15, encodes yet a third small RNA (Pfs1) that targets a different *C. elegans* gene, the Ephrin receptor gene *vab-1*, to induce avoidance and its transgenerational inheritance. While Pfs1 does not target *maco-1* itself, treatment of *C. elegans* with PF15 or reduction of *vab-1* through Pfs1 targeting results in the downregulation *maco-1,* suggesting that they act in the same pathway to regulate avoidance. This suggests that *C. elegans* has developed an ability to detect different small RNAs from multiple bacterial species. Like the PA14 and *P. vranovensis* transgenerational avoidance pathways, PF15 avoidance uses *Cer1* particles for germline-to-neuron signaling and DAF-7 signaling in the ASI neurons, indicating that the molecular mechanisms are conserved.

Intriguingly, RNAi knockdown of *vab-2*, which encodes an Ephrin ligand, also results in PF15 avoidance. The fact that VAB-1 and VAB-2 are involved in this response suggests an adult role for the Ephrin axon guidance pathway in a secondary adult function, regulation of bacterial avoidance. We recently found that several axon guidance genes play a role in the regulation of memory both in worms and in mice, suggesting a previously unappreciated role for these pathways in adult behavioral functions^30^. Thus, genes that are considered primarily “developmental” may in fact have secondary functions that only emerge in adulthood.

The fact that we have now identified three different small RNAs in three different *Pseudomonas* species that use three different short (16-17nt) perfect match sequences to target two *C. elegans* genes that act in the same pathway has several different implications. *C. elegans* appears to have evolved a pathogen avoidance strategy that involves “reading” a *Pseudomonas* pathogen’s small RNAs as biomarkers of future illness by matching perfect sequences of genes involved in the pathogen avoidance behavior, including *vab-1* and *maco-1*. The resulting adaptive immune memory provides a powerful survival advantage for subsequent progeny, allowing them to properly respond to a pathogenic threat without ever experiencing infection and illness, but still turn down this avoidance after four generations to prevent avoidance of beneficial food sources that “smell” like the pathogens^10^.

Thus, it is likely that other pathogenic *Pseudomonas* species in the worm’s environment also encode small RNAs that *C. elegans* interpret as biomarkers of future pathogenesis, perhaps targeting other genes in the *vab-1/maco-1/daf-7* avoidance pathway. The consistency of *C. elegans’* responses to small RNAs from *Pseudomonas* species found in the wild suggests that we have uncovered a robust and reproducible mechanism of transgenerational epigenetic inheritance that *C. elegans* has evolved to avoid pathogens without sacrificing nutritional resources.

## Methods

### Bacterial strains

P. fluorescens 15 (PF15) was a gift from M. Donia. E. coli OP50 was provided by the CGC. S. marcescens (ATCC 274) was provided by the ATCC. Control (L44440), vab-1, maco-1, vab-2, aat-4, aman-2, catp-8, chd-7, clec-245, cutl-18, dod-23, F42F12.3, fbxb-99, lam-1, let-502, nfx-1, pix-1, rbp-2, sdz-31, snx-14, T11G6.4, Y49E10.18, Y53G8AR.9, Y82E9BL.6, Y92H12BR.7, ZK1053.4 RNAi were obtained from the Ahringer RNAi library and sequences were verified. E. coli expressing the 600-nt PF15 intergenic region containing the 16-nt match to vab-1 was constructed by Gibson assembly. The intergenic region was cloned out of PF15 and ligated into pZE27GFP using PCR to linearize the plasmid, DpnI digest, and a single fragment Gibson assembly to ligate. The plasmid was transformed into MG1655 E. coli using a standard transformation protocol.

pZE27GFP fwd: GGTACCTTTGTCCTCTTTAATG
pZE27GFP rev: taagcttgatgggggatcc
Intergenic region fwd: GGGATCCCCCATCAAGCTTATAAGACTTTAACCGA AATTGTTAG
Intergenic region rev: TTAAAGAGGACAAAGGTACCTCTTCGTTGCCT TCCATGC

*E. coli* expressing the Pfs1 small RNA was constructed in a similar method, using primers specific to the Pfs1 region. Pfs1 was ligated into pZE27GFP using PCR to open the plasmid, DpnI digest, and a single fragment Gibson assembly to ligate. This plasmid was transformed into MG1655 *E. coli* using a standard transformation protocol.

Pfs1 fwd: GGGATCCCCCATCAAGCTTATATTGCCATAAACGACTTCCTCTG
Pfs1 rev: TTAAAGAGGACAAAGGTACCAATTGTTAGAGGTGATGCCCAAGT

Control bacteria for intergenic region and Pfs1 expressing bacteria consisted of plasmid pZE27GFP transformed into MG1655 *E. coli* using a standard transformation protocol.

### C. elegans strains

The following worm strains were provided by the C. elegans Genetics Center (CGC): N2 (wild type), KU25: *pmk-1(km25),* FK181: ksIs2 [*Pdaf-7p::gfp + rol-6(su1006)]*, CQ759: *maco-1(ok3165)*. CQ667: *Cer1(gk870313)*. CZ414: *vab-1(e699)*.

### Cultivation of bacterial strains

*E. coli* OP50, *P. fluorescens 15* (PF15), and *S. marcescens* were cultured overnight in Luria broth (10 g/l tryptone, 5 g/l yeast extract, 10 g/l NaCl in distilled water), shaking (250 rpm) at 37 °C. *E. coli* expressing the intergenic region or Pfs1 sRNA were grown in LB supplemented with 50 μg/mL Kanamycin. RNAi bacteria were supplemented with 100 μg/ml carbenicillin and 12.5 μg/ml tetracycline.

### General maintenance of *C. elegans* strains

Worm strains were maintained at 20°C on high growth medium (HG) plates (3 g/l NaCl, 20 g/l bacto-peptone, 30 g/l bacto-agar in distilled water, with 4 ml/l cholesterol (5 mg/ml in ethanol), 1 ml/l 1 M CaCl2, 1 ml/l 1 M MgSO4 and 25 ml/l 1 M potassium phosphate buffer (pH 6.0) added to molten agar after autoclaving) on *E. coli* OP50 using standard methods.

### Training plate preparation

Training plates were prepared by seeding 800 uL of bacteria onto NGM (3 g/L NaCl, 2.5 g/L Bacto-peptone, 17 g/L Bacto-agar in distilled water, with 1 mL/L cholesterol (5 mg/mL in ethanol), 1 mL/L 1M CaCl2, 1 mL/L 1M MgSO4, and 25 mL/L 1M potassium phosphate buffer (pH 6.0) added to molten agar after autoclaving) or HG plates.

Pathogenic bacteria such as PF15 or *S. marcescens* and control *E. coli* OP50 were prepared on NGM plates to avoid pathogen overgrowth, while sRNA producing MG1655 were prepared on HG plates (supplemented with 50 μg/mL Kanamycin). For sRNA training, 200 μl of OP50 was spotted in the center of a 6-cm NGM plate. Plates were stored at 25°C for 48hrs. RNAi bacteria were prepared on HG plates (supplemented with 1 mL/L 1M IPTG, and 1 mL/L 100 mg/mL carbenicillin) and kept at room temperature for 48 hrs. Plates were left to cool on a benchtop for 1 h to equilibrate to room temperature before the addition of worms.

### Bacterial choice assay plate preparation

Overnight bacterial cultures of PF15 were diluted in LB to an OD600 = 1, and standard cultures of undiluted OP50 were used. 25 uL of each bacterial suspension was spotted onto one side of a 6 cm NGM plate to make bacterial choice plates. These plates were incubated for 2 days at 25°C and allowed to equilibrate to room temperature for 1 hr before performing choice assay.

### Preparation of bacteria for RNA isolation

Bacterial lawns (plate) for RNA collection were cultured as previously described and grown for 2 days on plates at 25 °C (or 37°C for sequencing). 37°C liquid cultures for sequencing were pelleted for collection. Bacterial lawns were collected from the surface of NGM plates using a cell scraper. In brief, 1 ml of M9 buffer was applied to the surface of the bacterial lawn, and the bacterial suspension following scraping was transferred to a 15-ml conical tube. PF15 from 10 plates or OP50 from 15 plates was pooled in each tube and pelleted at 5,000g for 10 min at 4 °C. The supernatant was discarded, and the pellet was resuspended in 1 ml of Trizol LS for every 100 μl of bacterial pellet recovered. The pellet was resuspended by vortexing and subsequently frozen at −80 °C until RNA isolation.

### Bacterial small RNA isolation

To isolate RNA from bacterial pellets, Trizol lysates were incubated at 65 °C for 10 min with occasional vortexing. Debris was pelleted at 7,000*g* for 5 min at 4 °C. The supernatant was transferred to new tubes containing 1/5 the volume of chloroform. Samples were mixed thoroughly by inverting and centrifuged at 12,000*g* for 10 min at 4 °C. The aqueous phase was used at input for RNA purification using the mirVana miRNA isolation kit according to the manufacturer’s instructions for small RNA (<200 nt) isolation. Purified RNA was used immediately for sRNA training, or for sequencing frozen at −80 °C until further use.

### Worm preparation for training

Eggs from young adult hermaphrodite worms were obtained by bleaching and subsequently placed onto HG plates and incubated at 20 °C for 2 days. Synchronized L4 worms (48-52 h post bleaching) were used in all training experiments. For sRNA training, 100 μg of sRNA was added to the OP50 spot on the sRNA training plates. For RNAi, 200 μL of 0.1M IPTG was spotted onto plates and left to dry at room temperature before adding worms.

Synchronized L4 worms were washed off plates using M9 and left to pellet on the bench top for 2-3 min. Then, 5 μl of worms were placed onto sRNA-spotted training plates, 10 μl or 40 μl of worms were plated onto OP50 or pathogen-seeded training plates, and 20 uL of worms were placed onto RNAi plates or MG1655 sRNA expressing bacteria plates. Worms were incubated on training plates at 20 °C in separate containers for 24 h. After 24 h, worms were washed off plates using M9 and washed an additional 2-3 times to remove excess bacteria. Worms were tested in the aversive learning assay.

### Aversive learning assay

On the day of the assay, choice assay plates were left at room temperature for 1 h before use. To start the assay, 1 μl of 1 M sodium azide was spotted onto each respective bacteria spot to be used as a paralyzing agent during choice assay and preserve first bacterial choice. Worms were washed off training plates in 2 mL M9 into 1.5 mL tubes and allowed to pellet by gravity. Worms were washed 2-3 additional times in M9. Using a wide orifice pipet tip, 5 μl of worms (approximately 100-200 worms) were spotted at the bottom of the assay plate, midway between the bacterial lawns. Aversive learning assays were incubated at room temperature for 1 h before manually counting the number of worms on each bacterial spot. Plating a large number of worms (>200) on choice assay plates was avoided, because excess worms clump at bacterial spots making it difficult to distinguish worms, and high densities of worms can alter behavior. In experiments in which F1 and subsequent generations are used: day 1 worms from parental (P0) training were bleached and eggs were placed onto HG plates and left for 3 days at 20 °C. After 3 days, the F1 worms (Day 1-72 hours) were washed off HG plates with M9 at day 1. Some of the pooled worms were subjected to an aversive learning assay, and the remaining worms were bleached to obtain eggs, which were then placed onto HG plates left at 20 °C for 3 days and used to test the F2 generation. The same steps were followed for subsequent generations.

### Sequencing and annotating the *P. fluorescens 15* genome

Genomic DNA was extracted from an overnight PF15 culture using the Qiagen DNeasy Blood and Tissue kit for gram-negative bacteria. Nanopore sequencing was performed by MIGS (now SeqCenter). Annotation of the genome was performed using Prodigal gene prediction software^31^ and Standalone BLAST. BioProject PRJNA1114673

### Bacterial sRNA sequencing

Each sample of PF15 sRNA was tested for *C. elegans* behavior before sequencing. The size distribution of sRNA samples was examined on a Bioanalyzer 2100 using RNA 6000 Pico chip (Agilent Technologies). The sRNA sequencing protocol was similar to the protocol used in^10^. Around 300 ng of sRNA from each sample was first treated with RNA 5′ pyrophosphohydrolase (New England Biolabs) at 37 °C for 30 min, then converted to Illumina sequencing libraries using the PrepX RNA-seq library preparation protocol on the automated Apollo 324 NGS Library Prep System (Takara Bio). In brief, the treated RNA samples were ligated to two different adapters at each end, then reverse-transcribed to cDNA and amplified by PCR using different barcoded primers. The libraries were examined on Bioanalyzer DNA High Sensitivity chips (Agilent) for size distribution, quantified by Qubit fluorometer (Invitrogen), and then pooled at equal molar amount and sequenced on Illumina NovaSeq 6000 S Prime flowcell as single-end 122-nt reads. The pass-filter reads were used for further analysis. BioProject PRJNA1114673

### Bacterial small RNA sequencing data analysis

2 replicates of PF15 small RNA grown from 25 °C plates were sequenced, 5 replicates of PF15 small RNA grown from 37°C plates were sequenced, and 2 replicates of PF15 small RNA grown from 37 °C liquid were sequenced. Reads were mapped to the sequenced PF15 genome using RNA STAR^32^. Default settings were used for the RNA STAR mapping. The resulting BAM files were then used to identify putative sRNAs.

### Whole worm mRNA isolation, sequencing, and analysis

Adult worms (P0) trained as described on *E. coli* OP50 or PF15 were collected in M9 and washed several times to remove excess bacteria. Worm pellets were crushed in liquid nitrogen and transferred to an appropriate volume of Trizol LS (100 μl of worm pellet in 900 uL of Trizol). Total RNA was extracted from Trizol using chloroform extraction, isopropabol and ethanol precipitation, and cleanup using the RNeasy mini kit. mRNA libraries for directional RNA sequencing were prepared using the SMARTer Apollo System and were sequenced (paired-end) on the Illumina HiSeq 2000 platform.

### Rapid amplification of cDNA ends for Pfs1 sRNA identification

Three independent replicates of PF15 sRNA were isolated from PF15 grown on plates at 25°C as described above and purified using the mirVana miRNA isolation kit. RACE to determine bacterial sRNA 5’ and 3’ ends was performed as described^33^. In brief, RNA was treated with DNAseI, XRN-1 endoribonuclease, and RppH with phenol/chloroform extraction and ethanol precipitation between each enzymatic step. Small RNAs were then ligated with T4 ligase and cDNA was made using the Superscript III First-strand system with random hexamers. cDNA fragments were amplified with PCR primers specific to the internal region of Pfs1, gel purified and cloned into the pCR™4-TOPO™ vector before sequencing with M13F.

Pfs1 RACE PCR F: CGCGAGACGGACGACTGACG
Pfs1 RACE PCR R: TCGCGGTACATCCACCACAGAG

### Pfs1 structure prediction

A prediction of the secondary structure of the Pfs1 sRNA was made using the mFold tool on the UNAfold webserver^34^.

### Imaging and image analysis

All images were taken on a Nikon Eclipse Ti microscope or Nikon A1 R confocal microscope. Day 1 worms were prepared and treated as described in ‘worm preparation for training’. Worms were mounted on 2% agar pads on glass slides and immobilized using 1 mM levamisole.

Differential inference contrast (DIC) images of whole worms following OP50, or PF15 lawn or sRNA training, were imaged at 20x following preparation and training as described in “worm preparation for training”. For GFP-transgenic worms, Z-stack multi-channel (DIC and GFP) images were acquired at 20x (*irg-1p::*gfp) or 60X (*dafp7p::gfp*) magnification.

For *irg-1p::gfp* quantification, worms were prepared as described. Image analysis was done with Fiji, where ROIs were drawn around each animal in the field of vision and mean intensity values for all regions of interest were recorded and plotted.

For *daf-7p::gfp* quantification, worms were prepared as described. Images were acquired and maximum intensity projections of head neurons were built using Fiji for representative images. Quantification of mean fluorescent intensity was done using NIS Elements software. Average pixel intensity was measured in each worm by drawing a Bezier outline of the neuron cell body for 2 ASI head neurons. For imaging *daf-7p::gfp* in each generation, exposure times were adjusted to prevent oversaturation. For control and treatment groups imaged on the same day, the same exposure time and camera settings were used.

### *C. elegans* survival assay

PF15 was grown in liquid culture overnight (37°C) and diluted 1:2 to an OD = 1. 750 μL of diluted PF15 or OP50 (control) was spread to cover six 6-cm NGM plates. Plates were incubated for 2 days at 25°C to allow bacterial growth. Plates were equilibrated to 20°C before adding worms (84 hours post-bleach) to plates. Survival assays were performed at 20°C, counting assay plates every 6-9h. Every 48h, worms were moved onto new plates.

## Quantification of *maco-1* gene expression by qPCR

RNA was isolated from F2 progeny of Pfs1 or control trained P0 animals as described above for whole worm mRNA isolation. Total RNA was extracted using chloroform extraction, isopropanol and ethanol precipitation, and cleanup using the RNeasy mini kit. cDNA was made from 1 μg of RNA using the Superscript III First-strand system for RT-PCR. The extracted cDNA was used as input for qPCR reactions using the Power SYBR green qPCR master mix and protocol and run on a Viia7 Real-time PCR system as previously described^10^. qPCR primers:

maco-1 Forward: GTGTCACGACAATTGCC

maco-1 Reverse: CACATAGGTAGTGGCGAG

act-1 Forward^35^: GGCCCAATCCAAGAGAGGTATC

act-1 Reverse^35^: CAACACGAAGCTCATTGTAGAAGG

### Statistical analysis

Survival assays were assessed using Log-rank (Mantel-Cox) tests. For choice assays, populations of worms were raised together under identical conditions and were randomly distributed into treatment conditions. For all choice assays, each dot represents an individual choice assay plate (about 10–300 worms per plate). The box extends from the 25th to the 75 percentile, with whiskers from the minimum to the maximum values. For the comparison of choice indices between more than two genotypes, one-way ANOVA with Tukey’s multiple comparisons test was used. For comparisons of choice indices between genotypes and between conditions (naïve vs learned), two-way ANOVA with Tukey’s multiple comparisons test was used. Unpaired t tests were performed for comparisons between two groups. Experiments were repeated on separate days with separate populations, to confirm that results were reproducible. Prism 9 software was used for all statistical analyses.

## Acknowledgements

We thank M. Donia for sharing the PF15 bacterial strain, the *C. elegans* Genetics Center for worm strains; W. Wang, J. Arly Volmar, and J. Miller (Genomics Core Facility, Princeton University); the Murphy lab for discussion and feedback, and BioRender.com for model figure design software. The work was supported by an NIH Director’s Pioneer Award to CTM (NIGMS DP1GM119167), a Transformative R01 to CTM (1R01AT011963-01), a Ford Predoctoral Fellowship to RS, and an NIH training grant for support of RS, RB & RSM (NIGMS T32GM007388).

## Notes

### Competing Interest Statement

The authors have declared no competing interest.

